# Antifungal activity of silver nanoparticles during *in-vitro* culture of *Stevia rebaudiana* Bertoni

**DOI:** 10.1101/846733

**Authors:** Marco A. Ramírez-Mosqueda, Lino Sánchez-Segura, Sandra L. Hernández-Valladolid, Elohim Bello-Bello, Jericó J. Bello-Bello

## Abstract

Contamination by fungi and bacteria during the *in-vitro* propagation of plants leads to considerable losses of biological material and precludes phytosanitary certification. The anti-microbial effect of silver nanoparticles (AgNPs) may be an alternative for the eradication of *in-vitro* contaminants. This study evaluated the microbicidal activity of AgNPs on a recurrent fungus during the micropropagation of stevia (*Stevia rebaudiana* Bertoni). First, the fungus was isolated and identified at a molecular level by the sequencing and analysis of the ITS4/ITS5 rDNA region. The results of the phylogenetic analysis of various fungi species showed that the strain under study (16-166-H) belongs to the genus *Sordaria* and is 86.74% similar to *S. tomento-alba* (strain CBS 260.78). Subsequently, the inhibition of the growth of *S. tomento-alba* was tested under different concentrations of AgNPs (0, 25, 50, 100, and 200 mg L^−1^), observing that 50 and 100 mg L^−1^ achieve ca. 50% growth inhibition (IC_50_), while 200 mg L^−1^ produces a drastic inhibition. On the other hand, the shape and size of AgNPs was examined using transmission electron microscopy (TEM), and the transport and accumulation of AgNPs in *S. tomento-alba* cells were monitored through multiphoton microscopy. The morphological and fluorescence analyses showed that AgNPs display different sizes, with larger nanoparticles retained in fungal cell walls while smaller AgNPs penetrate into fungal cells. Probably, apoplastic and symplastic mechanisms involved in the accumulation and transport of AgNPs affect the metabolic processes of the fungus, thus inhibiting its growth. These results suggest that AgNPs possess antifungal activity and can be used in the eradication of contaminants during the *in-vitro* culture of plant species.

## Introduction

Plant tissue culture (PTC) is a biotechnological technique used for the *in-vitro* conservation, handling, sanitation, and propagation of edible, medicinal, and ornamental plants. *In-vitro* propagation, or micropropagation, represents a commercial alternative to produce pathogen-free plants (Efferth 2019). The success of micropropagation depends on ensuring strict asepsis. However, microbial contamination may occur during the micropropagation of plants, leading to important losses of plant material *in vitro* (Medjemem et al. 2016).

PTC contamination may be due to endophyte microorganisms, anthropogenic factors, tolerance of microorganisms to autoclaving, and resistance to antibiotics and fungicides (Thomas et al. 2017; Chechi et al. 2019; Marjon et al. 2019). *In-vitro* contaminants can affect the growth of explants by competing for water, light, space, and essential nutrients (Javed et al. 2017; Khan et al. 2018). In addition, the presence of contaminants limits the phytosanitary certification of PTC plant material (Sastry et al. 2014; Whattam et al. 2014). Phytosanitary certification is a priority issue in government policies in relation to economic income from the exportation and importation of *in vitro* plants of commercial interest (Whattam et al. 2014; Eschen et al. 2015).

There are several techniques for contamination control in micropropagated plants. One of them involves the treatment of parent plants and explants by adding fungicides and antibiotics to the culture medium (Cassells 2012). Nonetheless, the addition of antibiotics or fungicides to the culture medium for controlling bacterial contamination is not recommended due to the resistance of some strains (Caniça et al. 2019; Chechi et al. 2019). Biofilms with microbicidal effect are also available and can be used to prevent contamination, such as Plant Preservative Mixture^®^ (PPM) and Vitrofural^®^ (G1); however, their limited availability restrain their commercial application. An alternative for the eradication of *in-vitro* contaminants is the use of silver nanoparticles (AgNPs). These have been used as antimicrobial agents for the *in-vitro* culture of various plant species (Spinoso-Castillo et al. 2017; Tung et al. 2018). The mechanisms of action of AgNPs as antifungal agent have not been fully elucidated because most studies have addressed antiviral and antibacterial properties (Pařil et al. 2017; Khezerlou et al. 2018).

The culture of Stevia (*Stevia rebaudiana* Bertoni) is of high commercial value because of the non-caloric steviosides and rebaudiosides contained in its leaves (Debnath et al. 2019; Rouhani et al. 2019). The commercial propagules currently produced are insufficient given the low percentage of seed germination and the reduced number of cuttings that adapt to soil (Angelini et al. 2018). The *in-vitro* propagation of this species is affected by spontaneous contamination after its establishment and during subculture. For this reason, it is necessary to develop micropropagation systems for this species to ensure the production of plants that are free of diseases and pathogens. The objective of this study was the identification and control of contamination during the *in-vitro* establishment of *S. rebaudiana* using silver nanoparticles.

## Materials and Methods

### Physicochemical characterization of silver nanoparticles by transmission electron microscopy

The AgNPs used in this study, formulated as Argovit^®^, were provided by the Production Centre Vector-Vita Ltd, located in Novosibirsk, Russia. Argovit^®^ is made up of 12 mg mL^−1^ of metallic silver and 188 mg mL^−1^ of polyvinylpyrrolidone (PVP, 15-30 kD). The morphology of nanoparticles was examined under a Philips/FEI Morgagni M-268 transmission electron microscope (Brno, Czech Republic). For the morphological analysis, 5 μL of particles in suspension were mounted on a copper grid of Formvar 300 mesh/carbon (Electron Microscopy Science, PA). Samples were dried at room temperature for 5 min. The operating conditions in all experiments were: high voltage (EHT) of 80 kV, high magnification of 1000-140000X, and working pressure of 5 × 10^−3^ Pa (5 × 10 −5 Torr). Micrographs were captured in tagged image file (.tif) format with a resolution of 1376 × 1032 pixels and a grey scale. In this format, 0 was assigned to black and 255 to white in the grey scale.

### Plant Material

The explants used were nodal segments of stevia (*Stevia rebaudiana* Bertoni cv. Morita II) measuring 2 cm in length that contained one axillary bud. The explants were disinfected with a surfactant solution (Tween-20/distilled water) and washed with a slow flow of running water for 30 minutes. Subsequently, in a laminar flow hood, explants were immersed in 70% (v/v) ethanol for 30 s and in 0.6% and 0.3% (v/v) sodium hypochlorite for 10 and 5 min, respectively. Three rinses with sterile water were performed. Finally, the explants were transferred to test tubes containing MS medium (Murashige and Skoog 1962), supplemented with 1 mg L^−1^ BA (Bencilademina, Sigma-Aldrich, St. Louis, MO), 30 g L^−1^ sucrose, and 2.5 g L^−1^ Phytagel™ (Sigma-Aldrich, St. Louis, MO). The pH of the media was adjusted to 5.8 ± 0.2. The tubes with culture medium were autoclaved at 124 KPa for 15 min. Cultures were incubated at 25 ± 2 °C with a 16/8 h photoperiod (light/dark), under an irradiation of 40-50 μmol m^−2^ s^−1^ provided by fluorescent lamps. Subsequently, the explants showing evidence of contamination were isolated.

### Isolation and culture of the fungus Sordaria tomento-alba

Discs with mycelia from contaminated explants were transferred with a scalpel to Petri dishes containing potato dextrose agar medium (PDA) (Sigma-Aldrich, St. Louis, MO). Subsequently, these discs were incubated at 27 °C for 72 hours.

### Molecular identification of contaminating microorganisms

#### DNA extraction, PCR amplification, and ITS sequencing

Genomic DNA was extracted from fungal mycelium using an alkaline lysis method (Doyle and Doyle 1987). DNA quality was measured using a Nanodrop^®^ ND-1000 spectrophotometer (Thermo Scientific, Wilmington, USA). Polymerase chain reaction (PCR) was performed using universal internal transcribed spacers (ITSs): ITS4 (5’- TCCTCCGCTTATTGATATGC-3’) and ITS5 (5’- GGAAGTAAAAGTCGTAACAAGG-3’) primers (White *et al.* 1990). The PCR final volume of the reactions was 20 μl, containing 50 ng of genomic DNA, 1X of PCR buffer (Invitrogen, EU), 0.8 mM of dNTPs (Invitrogen, EU), 3 mM MgCl_2_, 0.5 μM of each primer and 1 U of *Taq* DNA Polymerase (Invitrogen, EU). DNA amplification was performed in a GeneAmp^®^ PCR System 9700 thermal cycler (Perkin-Elmer). PCR parameters consisted of one cycle of initial denaturation at 90 °C for 30 s, followed by 35 cycles of denaturation at 90 °C for 15 s, primer annealing at 56 °C for 30 s, elongation at 72 °C for 1 min, and a final elongation at 72 °C for 7 min. The amplification products were separated by electrophoresis in 1.2% (w/v) agarose gel previously stained with ethidium bromide. The run was performed in TAE 1X buffer (Tris-Acetic acid-EDTA) at 100 V for 45 min. PCR products were sequenced at the Colegio de Postgraduados (Campus Montecillo) using the HiSeq 2500® Sequencing System – Illumina (Sanger method). PCR products were sequenced at the Colegio de Postgraduados (Campus Montecillo) using the HiSeq 2500^®^ Sequencing System – Illumina (Sanger method).

#### Multiple sequence alignment and phylogenetic tree construction

ITS4/ITS5 rDNA from the 16-166-H strain was used as query sequence and compared against the NCBI database using the BLAST nucleotide search tool (https://blast.ncbi.nlm.nih.gov/Blast.cgi) (Altschul *et al.*, 1997). An *in-silico* analysis was developed with 15 target sequences producing significant alignments, using the *Colletotrichum acutatum* (MH865675) sequence as outgroup control. The multiple sequence alignment was performed using the ClustalW algorithm in the msa package (version 1.16.0) from the R program (Bodenhofer et al. 2015). For the phylogenetic analysis, the ape package (version 5.3) was employed (Paradis and Schliep 2019). The phylogenetic relations of samples were constructed using the Neighbor-Joining method, and the genetic distances were computed using the Jukes-Cantor method (Jukes and Cantor 1969; Saitou and Nei 1987). The optimal tree was generated with 1000 bootstrap replicates. Bootstrap support threshold equal or greater than 50% was considered significant. Graphical tree representation was plotted with the ggtree package (Yu et al 2017).

#### Inhibition of fungal growth

The antimicrobial activity of AgNPs on the growth of *S. tomento-alba* was explored using mycelia seeded in plates with PDA and evaluating different concentrations of AgNPs (0, 25, 50, 100, and 200 mg L^−1^). First, the culture medium was adjusted to a pH of 6.5 and was sterilized at 124 KPa por 15 min. Then, all treatments were inoculated with 1 cm^2^ of fresh mycelium of *S. tomento-alba* and incubated under a photoperiod of 18/6 hours of light/darkness, at 23-25 °C. After 5 days of incubation, the variable to measure was fungus growth in diameter (known as GD), with the average of three measurements (in cm) considered as GD. The growth of *S. tomento-alba* was evaluated after 21 days of incubation.

For the fungus growth in diameter (cm), a completely randomized design was used, as described below:

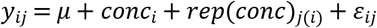

Where *y*_*ij*_ is fungus diameter observed at concentration *i* of AgNPs in replicate *j*; μ is the overall mean; *conc*_*i*_ is the fixed effect of concentration *i* of AgNPs; *rep*(*conc*)_*j*(*i*)_ is the random effect of replicate *j* nested on concentration *i* of AgNPs assuming 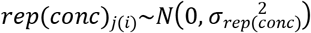 and *ε*_*ij*_ is the experimental error with *ε*_*ij*_~*N*(0,σ^2^). Fungus growth was analyzed with the procedure PROC GLIMMIX of SAS (version 9.4) under a Generalized Linear Mixed Model with Poisson distribution, and for fungus growth, Linear Mixed model was used.

#### Detection and action of AgNPs in fungal inhibition by fluorescence microscopy

AgNPs were detected using modifications of the “Lambda” method for determining the native spectral emission of AgNPs (Castro-González et al. 2019). The nanoparticles accumulated in hyphae were visualized with a multiphoton microscopy system (Axio Imager Z2, LSM 880-NLO, Zeiss, Oberkochen, Germany) coupled to a Ti: Sapphire infrared laser (Chameleon Vision II, COHERENT, Santa Clara, CA, USA) with a tuning capability in the range of 690 to 1060 nm. In all experiments, the operating conditions involved the use of a Chameleon laser set at 850 nm with 1.5% power, pinole at 600.1, and similar photodetector voltage ranges. Emissions from AgNPs were recovered at 596–637 nm. Images of hyphae were captured with a 63X/1.40 immersion objective, and NA ∞-0.17, using a Zeiss Plan NEOFLUAR with a 5 nm spectral sensitivity. All micrographs were captured in CZI format in a size of 1131×1131 pixels composed of three color channels (RGB).

## Results

### Physicochemical characteristics of Argovit^®^

The physicochemical characteristics of AgNPs are shown in Table 1. The AgNPs characterized by TEM are spherical with a form factor of (0.82) and roundness of 0.88. The analysis of AgNP dimensions showed average diameters of 35 ±15 nm, which consists of clustered silver (12 mg/mL metallic silver) functionalized with 188 mg/mL of polyvinylpyrrolidone (PVP, 10-30 kD). The results obtained evidence the structural dimensions of AgNPs (Argovit^®^-CP) in terms of shape and size. The size of AgNPs was verified by TEM, showing macroscopic aggregates composed of silver nanoparticles. The TEM micrograph (Figure 1) corroborates the tendency to aggregate; NPs of different sizes were observed, showing spheroidal nanoparticles ranging from 13 nm to 80 nm.

**Table 1.** Physicochemical characteristics of Argovit^®^

| Properties | Mean |
| --- | --- |
| Average diameter of metallic silver particles by TEM (nm) | 40±10 |
| Form Factor (Spheroid) | 0.80 |
| Metallic silver content (% wt.) | 1.2 |
| PVP Content (% wt.) | 18.8 |
| Roundness | 0.80 |
| Size interval of metallic silver particles by TEM (nm) | 2 - 85 |
| Zeta potential (mV) | -15 |

Abbreviations: Ag, silver; PVP, polyvinylpyrrolidone; TEM, transmission electron microscopy.

**Fig. 1.**
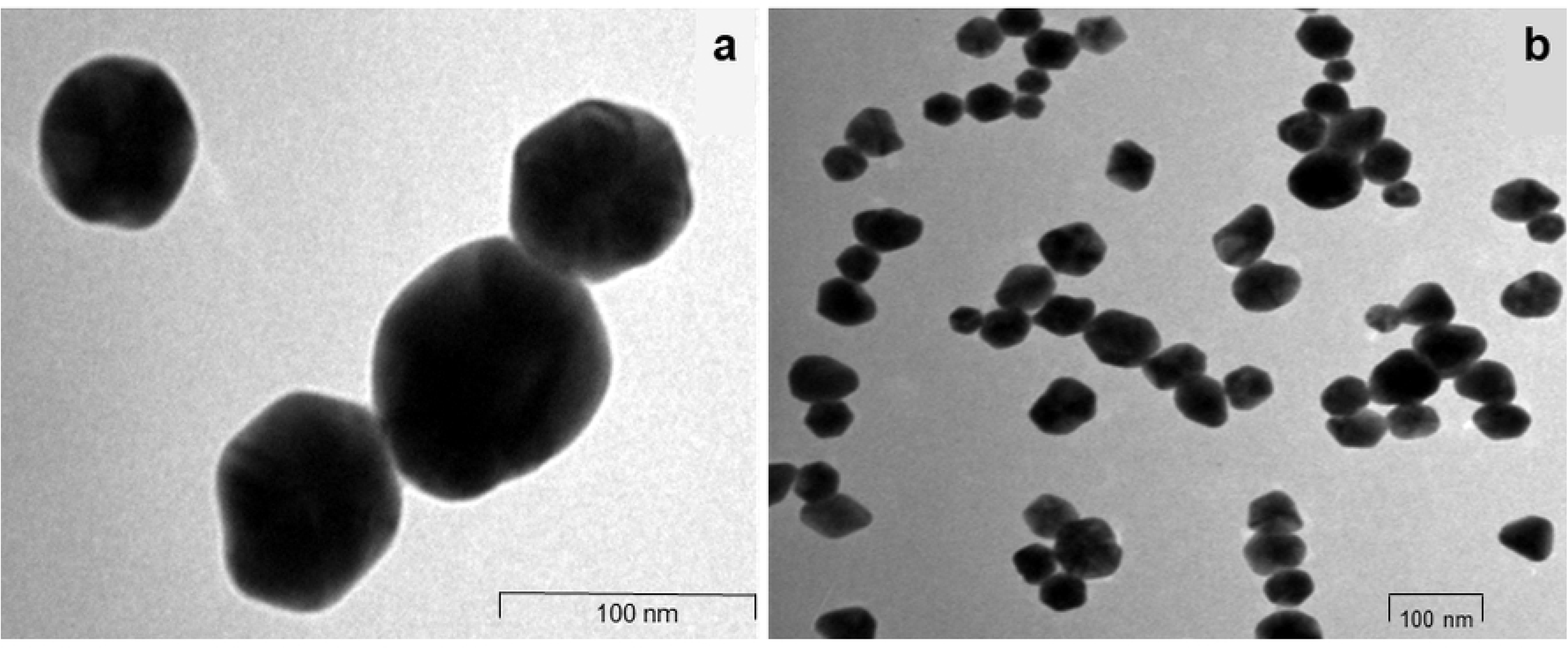
Microphotographs of Argovit® AgNPs. a) Magnified TEM images of an AgNP aggregate of spherical shape and several sizes in the range of 36.59-66.24 nm. Scale bar=100 nm. b) TEM micrograph of AgNP aggregates. Scale bar = 500 nm.

### Molecular identification of the 16-166-H fungal strain

The BLAST algorithm-based analysis of the 560-bp ITS sequence showed that this fungal strain had a high percent identity value (96.64%) to *Sordaria* strains JN207345, JN207271, and JN207268, which are associated to *Sordaria tomento-alba* as reported by Loro et al. 2012. Moreover, these results were confirmed by the phylogenetic tree analysis. The topology of the phylogenetic tree showed the formation of 5 clades. Among these clades, *Sordaria* and *Asordaria* species were grouped in three internal clades (Fig. 2). Clade I comprises *Sordariomycetes* sp., *Sordaria fimicola,* and *Sordaria fimicola*; Clade II comprises *Sordaria tomento-alba* and related *Sordaria* strains (including 16-166-H); finally, Clade III comprises *Sordaria* sp., *Asordaria prolifica*, and *Asordaria conoidea*. Both results reveal insights into the molecular identification of the fungal isolate. For this analysis, a reference strain was deposited in Genbank (https://www.ncbi.nlm.nih.gov/genbank/).

**Fig. 2.**
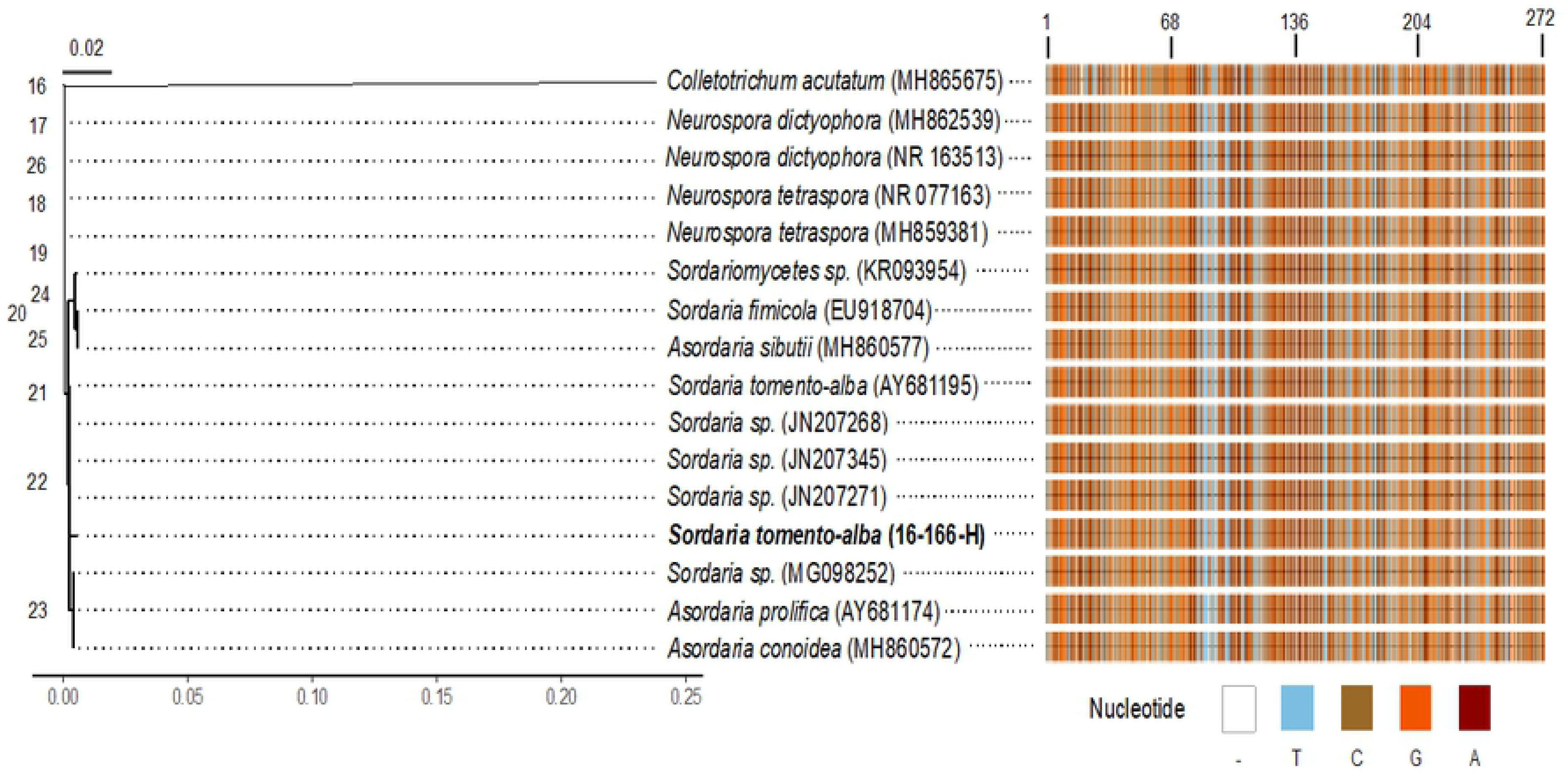
Phylogenetic tree and multiple sequence alignment of 16-166-H strain based on ITS4/ITS5 rDNA and NCBI BLAST sequences. A) The phylogenetic tree was inferred from a distance analysis with the Neighbor-Joining method. Colletotrichum acutatum (MH865675) was used as outgroup. B) An abstract multiple sequence alignment of 16 NCBI BLAST sequences was performed with the ClustalW algorithm. Numbers at the top of the graph correspond to columns in the alignment (bp). Nucleotide legends are shown at the bottom.

### Inhibition of fungal growth

The results show the fungicidal effect of AgNPs on the growth of *S. tomento-alba* in solid culture medium. AgNP concentrations of at least 50 mg L-1 produced a significant inhibition on fungal growth (Fig. 3). The highest greatest fungal GD was observed at 0 and 25 mg L^−1^ NPsAg, with 7.00 ± 0.54 and 7.60 ± 0.34 cm in diameter, respectively, whereas the lowest occurred at 200 mg L^−1^, with 2.50 ± 0.05 cm (Fig. 4). For 50 and 100 mg L^−1^, no significant differences were observed in the development of the fungus, recording an average diameter of 4.30 ± 0.60 and 4.13 ± 0.41 cm, respectively. (Fig. 3).

**Fig. 3.**
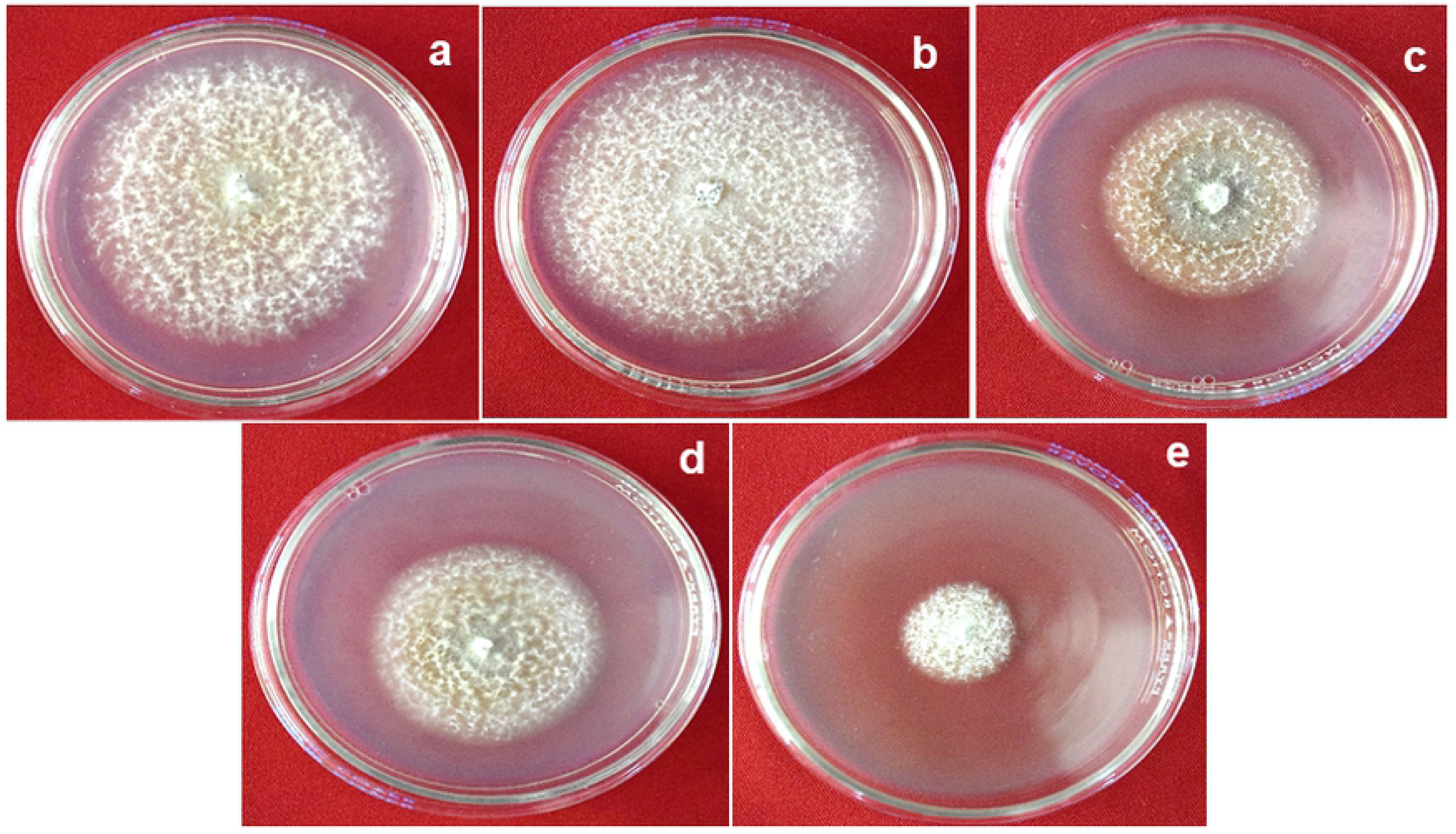
Effect of different AgNP concentrations on growth of Sordaria tomento-alba in PDA medium after 21 days. Bars represent the mean ± standard error. Bars followed by different letters denote significant statistical differences (Tukey, p ≤0.05).

**Fig. 4.**
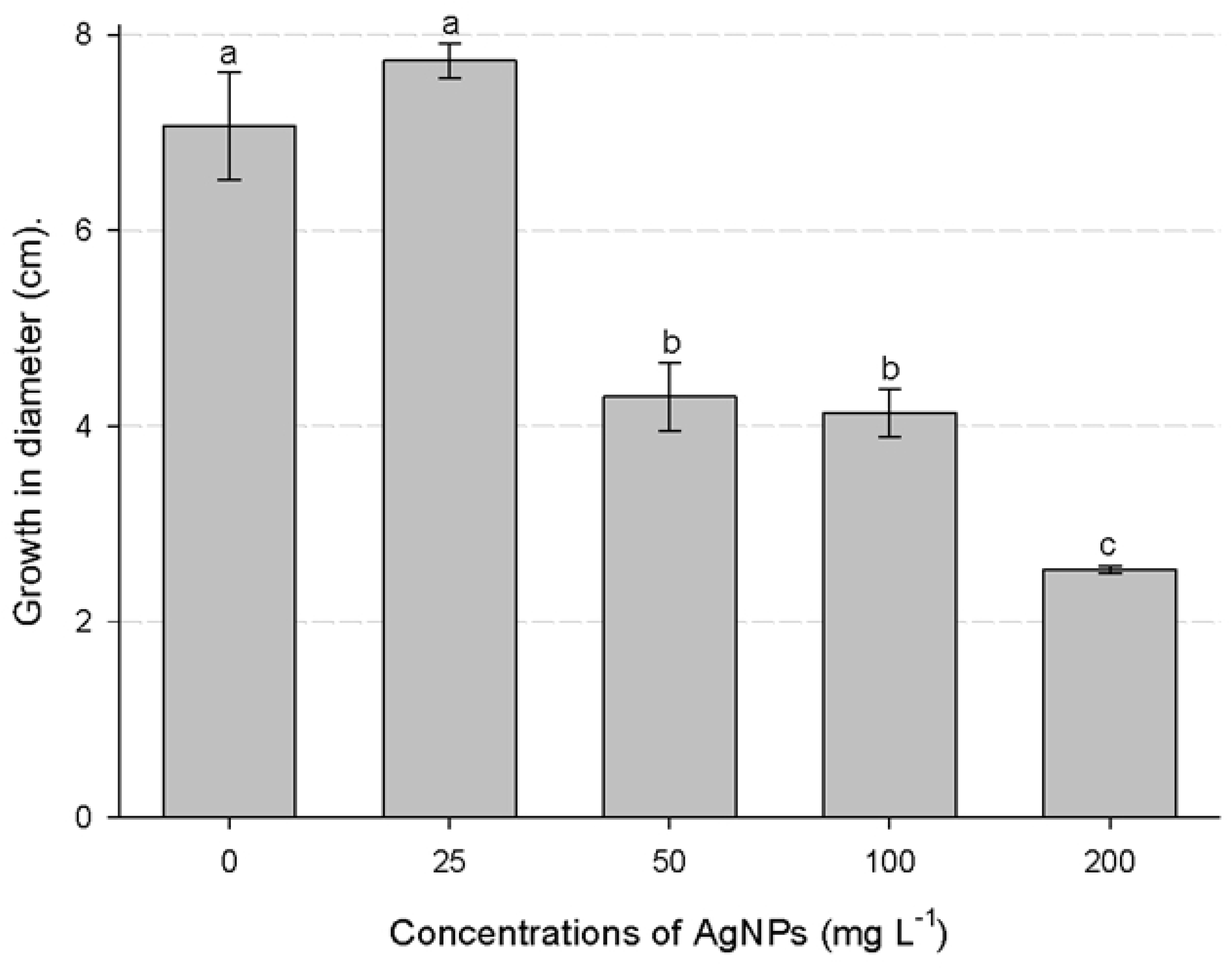
Growth of S. tomento-alba in PDA supplemented with different concentrations of AgNPs after 21 days of incubation: a) 0 mg L-1, b) 25 mg L-1, c) 50 mg L-1, d) 100 mg L-1, and e) 200 mg L-1.

### Detection and effect of AgNPs on fungal growth inhibition by fluorescence microscopy

Multiphoton microscopy allowed observing the presence of AgNPs in fungus cells. It evidenced the presence of AgNPs in the cell wall of hyphae (cross-section) subjected to the different treatments with NPs. However, as AgNP concentration increased, nanoparticles showed a trend to accumulate in the space between the cell wall and the cell membrane. The sequence of images in Fig. 5 shows the clear field and fluorescence of AgNPs and the progression of fluorescence in stem cross-sections under different treatments with AgNPs.

**Fig. 5.**
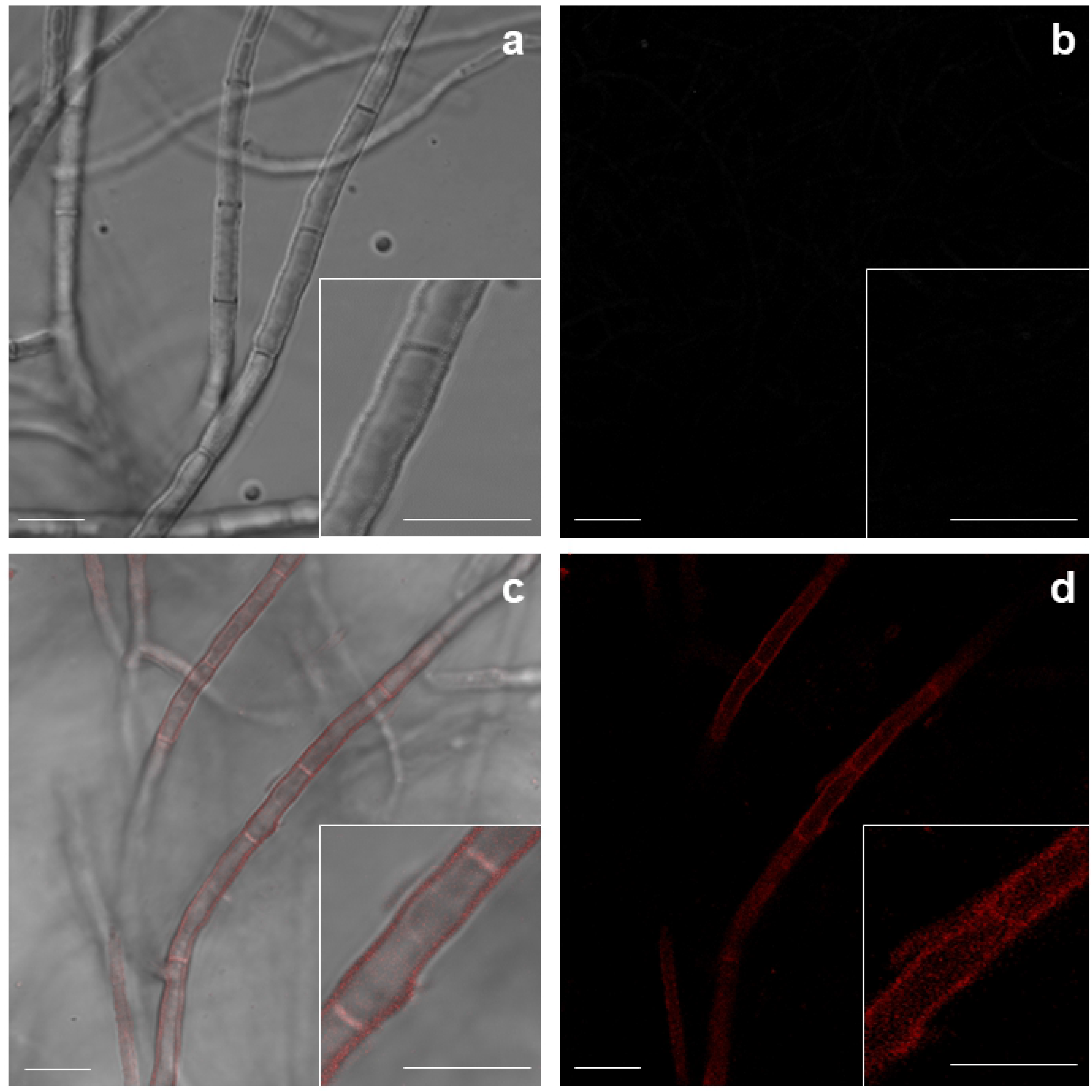
Location of AgNPs in S. tomento-alba cultured in PDA media. a-b) 0 mg L-1 AgNPs, merging and fluorescence, respectively, c-b) 200 mg L-1 AgNPs, merging and fluorescence, respectively. Bar = 10 μm.

## Discussion

This study evidenced the antifungal effect of AgNPs on the fungus *S. tomento-alba* during the *in-vitro* establishment of *S. rebaudiana*. *S. tomento-alba* has been reported as an endophyte in various plant species, including *Stryphnodendron adstringens* (Carvalho et al. 2012), *Cenchrus ciliaris*, and *Cenchrus* cf. *spinifex* Cav. (Loro et al. 2012), and in different cultivars of *Solanum tuberosum* (Zimudzi et al. 2017). Although this fungus is not reported as a phytopathogen, it causes contamination issues in *in-vitro Stevia* cultures. The effectiveness of AgNPs in the elimination of microbial contaminants from *in-vitro* cultures depends on AgNP size, shape, and type of coating. With regard to the characterization of Argovit^®^, the TEM allowed us to observe particles of 40 ± 10 nm in size, with the dominance of a spheroid form factor (0.80). The toxicity of AgNPs in biological systems has been reported to be inversely proportional to particle size (i.e., smaller particles are more toxic) (Panzarini et al. 2018).

The analysis of the ITS sequence of the fungal strain 16-166-H supported its identification as belonging to the genus *Sordaria*, genetically related to the species *S. tomento-alba.* The molecular identification of contaminants is relevant because it allows knowing its origin and deriving an appropriate treatment for the disinfection of explants (Tomasi et al. 2017). In this study, it was found that *S. tomento-alba* is a non-phytopathogenic endophyte. However, it can lead to contamination in the *in-vitro* establishment of *S. rebaudiana* and, if uncontrolled, may spread across the laboratory and affect other species cultured *in vitro*. The AgNPs used in this study probably affect the growth of species of the genus *Sordaria* (*Sordaria* spp.). In general, *in-vitro* contaminants first use the carbon available within the plant to survive; subsequently, they migrate to the culture medium that is rich in nutrients and contains sucrose as carbon source. The fungi in the culture medium compete with explants for space, water, light, and nutrients, causing the death of explants (Cassells 2012; Tomasi et al. 2017). One option to eradicate these microorganisms is through the culture of meristems, thermotherapy, and use of antifungal agents (Cassells 2012; Smith 2013: Sasi and Bhat 2018). The culture of meristems has been used primarily to eradicate viruses (Sasi and Bhat, 2018); for its part, thermotherapy occasionally damages the explants (Hu et al. 2019; Kaur et al. 2019). As regards the use of antifungal agents, these can lead to the development of resistance in some fungal strains (Caniça et al. 2019; Chechi et al. 2019). AgNPs do not exert selective pressure on microorganisms and, therefore, do not lead to resistance (Lemire et al. 2013; Khezerlou et al. 2018); thus, these may be less toxic than synthetic fungicides.

In our study, during the inhibition of the growth of *S. tomento-alba,* we noted that AgNP concentrations of 50 and 100 mg L^−1^ correspond to the half maximal inhibitory concentration (IC_50_). An antifungal effect of AgNPs has been reported for different fungi strains. Jo et al. (2009) mention that AgNP concentrations from 200 mg L^−1^ were needed to control the development of spores of *Bipolaris sorokiniana* and *Magnaporthe grisea*. On the other hand, Pulit et al. (2013) demonstrated the inhibition of the growth of *Cladosporium cladosporioides* and *Aspergillus niger* strains at 50 mg L^−1^ AgNPs. Kim et al. (2009) observed that 25 mg L^−1^ of AgNPs affect the integrity of the structure of hyphae of *Raffaelea* spp., while Kasprowicz et al., (2010) noted that 10 mg L^−1^ of AgNPs reduce the radial growth of *Fusarium culmorum*. Moreover, AgNP concentrations from 2.5 mg L^−1^ drastically reduce the germination of spores. Recently, Ruiz-Romero et al. (2018) used AgNPs to inhibit the radial growth of two phytopathogenic fungi (*Fusarium solani* and *Macrophomina phaseolina*). However, this work only mentions that the AgNPs supplemented were obtained as an extract from Yucca (*Yucca shilerifera*), without reporting the AgNP concentration. Also, the physicochemical characteristics of AgNPs are not reported in the studies mentioned above. It is our opinion that AgNPs should be characterized before conducting a research study.

The microbicidal effect of AgNPs derives from the interaction of silver ions with a broad range of molecular and metabolic processes within organisms, including growth inhibition, cell death, and inhibition of DNA replication (Abdi et al. 2008; Yun’an Qing et al. 2018). In fungi, AgNPs break the cell membrane of hyphae, thus impairing infection mechanisms (Kim et al. 2008; Bocate et al. 2019) and inhibiting the germination of conidia (Kim et al. 2009). In our study, we observed the accumulation of AgNPs in the cell wall and cytoplasm of hyphae of *S. tomento-alba*. The exact mechanism of transport and accumulation of AgNPs in fungi is currently unknown. However, the results of the characterization of AgNPs suggest that due to diversity of sizes (2-85 nm), larger nanoparticles accumulate in the cell wall and cell membrane, affecting the integrity of these organelles; for their part, smaller nanoparticles penetrate through pores in the cell wall and the cell membrane via transport by plasmodesmata. According to Money (1990), cell membrane pore size in hyphae of some fungi ranges from 2.3-3.3 nm, which could explain the penetration of small nanoparticles. AgNPs that manage to penetrate inside the cell cause an increase of Ag+ cations, which could affect the electrical potential of the membrane. According to Srikar et al. (2016) and Khezerlou et al. (2018), these Ag+ ions denaturate proteins, deplete intracellular ATP, and form complexes with DNA bases, so that the DNA loses its replication ability. On the other hand, the reaction of Ag + with thiol, carboxylate, phosphate, hydroxyl, amine, imidazole, and indole groups in enzymes may lead to their inactivation and cell death (Lin et al. 1998; Ashraf et al. 2013). The Ag+ in nanoparticles probably exerts important effects on biological systems. The use of AgNPs for disease control has the advantage of being non-toxic to humans and the environment, unlike synthetic pesticides.

In conclusion, the AgNPs used in this study showed an antifungal effect in *S. tomento-alba*, a common contaminant during the establishment of *S. rebaudiana*. AgNPs with similar physicochemical characteristics may be used to control other fungal strains that contaminate *in-vitro* cultures. Therefore, further studies should be conducted on the microbicidal potential of AgNPs in the micropropagation of different plant species, and the effects of AgNPs on DNA damage and replication.

## Author contribution statement

SLH and JJBB devised and designed this research. SLH and LSS conducted the experiments. MARM and EBB performed and reviewed the molecular and statistical analyses. MARM and JJBB drafted the manuscript. All authors reviewed and approved the manuscript.

